# Multiple viral protein genome-linked proteins compensate viral translation in a +ssRNA virus infection

**DOI:** 10.1101/2022.05.27.493807

**Authors:** Reid Warsaba, Nikolay Stoynov, Kyung-Mee Moon, Stephane Flibotte, Leonard Foster, Eric Jan

## Abstract

Viral protein genome-linked (VPg) protein plays an essential role in protein-primed replication of plus stranded RNA viruses. VPg is covalently linked to the 5’ end of the viral RNA genome via a phosphodiester bond typically at a conserved amino acid. Whereas most viruses have a single VPg, some viruses encode multiple VPgs that are proposed to have redundant yet undefined roles in viral replication. Here, we use the dicistrovirus, cricket paralysis virus (CrPV), which encodes four non-identical copies of VPg, as a model to characterize the role of VPg copies in infection. Dicistroviruses encode two main open reading frames (ORFs) that are driven by distinct IRESs. We systematically generated single and combinatorial deletions and mutations of VPg1-4 within the CrPV infectious clone and monitored viral yield in Drosophila S2 cells. Deletion of one to three VPg copies progressively decreased viral yield and delayed viral replication, suggesting a threshold number of VPgs for productive infection. Mass spectrometry analysis of CrPV VPg-linked RNAs revealed viral RNA linkage to either a serine or threonine in VPg, from which mutations in all VPgs attenuated infection. Mutating serine 4 in a single VPg abolished viral infection, indicating a dominant-negative effect. Using viral minigenome reporters that monitor dicistrovirus 5’ untranslated (UTR) and intergenic internal ribosome entry site (IRES) translation revealed a relationship between VPg copy number and the ratio of distinct IRES translation. We uncover a novel viral strategy whereby VPg copies in dicistrovirus genomes compensate for the relative IRES translation efficiencies to promote infection.

**Importance:** Genetic duplication is exceedingly rare in small RNA viral genomes as there is selective pressure to prevent RNA genomes from expanding. However, some small RNA viruses encode multiple copies of a viral protein, most notably an unusual viral protein that is linked to the viral RNA genome. Here, we investigate a family of viruses that contains multiple viral protein genome-linked proteins and reveal a novel viral strategy whereby viral protein copy number counterbalances differences in viral protein synthesis mechanisms.

## INTRODUCTION

Viral protein genome-linked (VPg) is a small virally-encoded protein that has multifunctional roles in specific steps of some +ssRNA viral lifecycles including viral replication and translation. VPgs are encoded in +ssRNA viral genomes including animal, plant, and fungal RNA viruses and their functions have been primarily studied in picornaviruses, caliciviruses, and potyviruses (1, 2). VPg is covalently linked to the 5’ end of the viral RNA and serves as a primer for both positive- and negative-strand RNA synthesis (3). For calicivirus and potyvirus, the VPg is also linked to subgenomic viral RNAs (4). The VPg is linked to the viral RNA via a hydroxyl-containing phosphodiester bond to an acceptor residue, serine, threonine, or tyrosine (5). In picornaviruses, the VPg protein is uridylated at a conserved tyrosine residue by a protein complex containing the viral RNA dependent RNA polymerase (RdRp). Uridylatation of the VPg by the RdRP-containing complex is mediated by an adenosine-rich motif (AAACA in poliovirus) that acts as a template within the apical loop of a stem-loop RNA structure called the *cis* replication element (CRE) (6). The location of CRE is conserved within the 2C domain of the picornavirus genome (7). However, the CRE location is position-independent (8). Uridylation occurs through a “slide-back” mechanism over the conserved adenosine-rich motif of the CRE thereby leading to the addition of two uridines linked to tyrosine 3 of VPg (VPgpUpU) (9). Following this, VPgpUpU and RdRp are recruited to the negative-sense RNA to prime and synthesize the new positive-sense viral genome resulting in a covalently attached 5’ VPg positive-sense RNA genome (10). The 5’ VPg-linked viral genomes are then packaged in virions (11). The negative-sense RNA is also bound to VPg at its 5’ end through a proposed alternate priming mechanism (12, 13). How VPgs are recruited from the CRE to the 5’ ends of positive- and negative-strand viral RNAs is not completely understood.

VPg linkage can be regulated during the viral life cycle. In the early stages of picornavirus infections, the VPg is unlinked from the viral RNA genome by DNA repair enzyme, 5′-tyrosyl–DNA phosphodiesterase-2 (TDP2), to allow for viral protein synthesis, thus providing a viral strategy to separate viral translation and replication (14, 15). However, the relevance of VPg unlinkase from the genome is not fully understood as a VPg-RNA that cannot be unlinked using a chemically non-cleavable bond did not affect picornavirus translation or replication (16). VPg can also stimulate the poliovirus RdRP activity and act as an RNA chaperone (17, 18).

Besides mediating replication, the VPg is associated with other functions (19). In caliciviruses, sapoviruses, vesiviruses and noroviruses, VPgs are essential for cap-independent translation (20). The calicivirus VPg interacts directly with the cap binding protein eIF4E to initiate translation (20). These viruses are insensitive to the addition of cap analogues, suggesting that the binding interface of eIF4E with VPg differs from that of a canonical cap (21). In marine norovirus, VPg interacts directly with the eIF4G HEAT-1 domain instead of eIF4E to recruit the translational initiation machinery to the viral RNA (22). Finally, the VPg of potyvirus can act as a viral suppressor of RNA silencing by degrading an RNA silencing factor in plants (23) and VPg can act as a switch for encapsidation in picornaviruses (24).

Although most viruses encode a single VPg in their genome, the Foot and mouth disease virus (FMDV) genome encodes three consecutive non-identical VPgs (25). To our knowledge, FMDV is the only virus in the aphthovirus family to encode multiple VPgs. All three FMDV VPgs can be uridylated *in vitro* by the FMDV RdRP where VPg3 is most efficiently uridylated *in vitro* (26, 27). Multiple copies of VPg in FMDV confer a competitive advantage (25, 28). VPg3 is essential for maximal FMDV infection and the identity of the C-terminal residues of VPg3 is necessary for proteolytic processing between VPg3 and the downstream 3C viral protein (29). Moreover, it has been reported that FMDV VPg can block RIG-I-mediated immune signaling (30). Although extensively studied in FMDV, it is not clear if the role of VPg copies in FMDV can be extrapolated to other viral families encoding multiple VPgs.

Dicistroviruses are positive-sense single-stranded RNA viruses that primarily infect arthropods that are of economic and agricultural importance (31, 32). Infections by Taura syndrome virus (TSV) and honeybee dicistroviruses, such as Israeli acute paralysis virus (IAPV) can lead to losses of shrimp and honeybee populations, respectively. Dicistroviruses are indirectly linked to human health such as Triatoma virus which infects T. cruzi, the causative agent of Chagas disease. Members of the dicistrovirus family also include the Cricket paralysis virus (CrPV), Drosophila C virus (DCV), and Rhopalosiphum padi virus (RhPV) that have been models for studying insect innate immunity and viral translational control strategies. Dicistroviruses contain an ∼ 9 kb RNA genome encoding two main open reading frames (ORFs), ORF1 and ORF2. ORF1 encodes the nonstructural proteins such as a multifunctional viral RNAi suppressor, RNA helicase, viral 3C-like protease, and RNA-dependent RNA polymerase (RdRP) whereas ORF2 encodes the structural capsid proteins. In dicistrovirus infections, ORF2 is expressed in supramolar excess compared to ORF1 proteins (33–36). Translation of the two ORFs is driven by distinct internal ribosome entry site (IRES) mechanisms within the 5’untranslated region (5’ UTR) and intergenic region (IGR) IRES. Extensive biochemical and structural studies have shown that the IGR IRES uses a triple pseudoknotted RNA structure to mediate a factorless recruitment strategy to directly recruit the ribosome and initiate translation of ORF2 (35, 37–40). The 5’ UTR IRES has only been investigated in a limited number of dicistroviruses (41–43). Unlike the IGR IRES factorless mechanism, the CrPV and RhPV 5’ UTR IRESs require a subset of translation factors to direct IRES translation (41, 44, 45). The distinct IRES mechanisms in dicistroviruses allow for differential and temporal expression of ORF1 and ORF2 for optimal virus infection (46).

Dicistrovirus RNA genomes are VPg-linked and contain a 3’ poly A tail (47). Interestingly, the dicistrovirus genomes contain multiple consecutive VPgs encoded in ORF1 ranging between two to six VPgs with VPg lengths ranging between 13 to 19 amino acids (48). Edman degradation of the VPg proteins from purified virions of the dicistrovirus Plautia stali intestine virus (PSIV) revealed that a glutamic acid at the C-terminus demarcates the boundary for all three VPgs are subsequently processed (48). Alignment of the limited dicistrovirus VPg sequences did not reveal an obvious conserved amino acid between dicistroviruses that may serve as the VPg linkage, however, a serine, threonine, or tyrosine is present in all VPgs that may be the virally linked amino acid (48). The role and relevance of the VPg copies in dicistrovirus infection have not been yet investigated.

Using CrPV, we examined the role of its four tandem VPgs and identified the amino acid that is linked to the viral RNA. Deletion of VPgs in CrPV revealed a minimum threshold for productive infection. Mass spectrometry analysis revealed that the CrPV viral RNA is linked to the serine at position 4 (S4) or threonine at position 9 (T9) of VPg. Mutational analysis of the 4^th^ serine or 9^th^ threonine of all VPgs abolished CrPV infection, demonstrating that both residues are necessary for CrPV infection. Mutating the serine 4 in a single VPg also inhibited virus infection, suggesting a dominant negative effect which was not observed following the mutation of a single threonine 9 nor deletion of a single VPg protein. To our knowledge, this is the first report of uridylation on two residues on a VPg. Finally, we demonstrate that the relative 5’ UTR IRES and IGR IRES translation efficiencies are directly related to viral genome VPg copy number, thereby revealing a viral gene duplication strategy to compensate viral translation mechanisms.

## RESULTS

### Threshold Number of VPg for Productive CrPV Infection

The CrPV genome contains four VPg copies, each of which is 18 amino acids in length with several identical residues including valine and glutamic acid at positions 1 and 18 (Figure 1A). We first addressed whether a minimal number of VPgs is necessary for CrPV infection. To test this, we generated single or consecutive VPg deletions within the CrPV infectious clone (Figure 1A) (49). Wild-type (WT) and deletion mutant CrPV RNAs were transfected into Drosophila S2 cells, followed by immunoblotting for the CrPV structural protein VP2 at 72 hours post-transfection (h.p.t.). In general, detection of VP2 at 72 h.p.t. is indicative of productive virus replication (49). As a negative control, we also transfected a CrPV(ORF1-STOP) RNA which contains a stop codon at the N-terminus of ORF1 effectively blocking CrPV translation and replication (49). As expected, the wild-type but not the CrPV(ORF1-STOP) mutant produced robust VP2 expression at 72 h.p.t. (Figure 1B). Deletion of a single VPg (ΔVPg1) or two VPgs (ΔVPg1-2) resulted in VP2 production whereas deletion of three consecutive VPgs (ΔVPg1-3) abolished VP2 production (Figure 1B), thus suggesting a minimal threshold of two VPgs for CrPV infection. To further support this, we monitored viral yield from transfected cells. The ΔVPg1 CrPV clone resulted in similar titres to wild-type CrPV 72 h.p.t., whereas ΔVPg1-2 resulted in significantly reduced titres and deletion of three VPgs abolished viral yield, which is in line with the immunoblotting analysis (Figure 1B, 1C). To further investigate the level of replication, we monitored CrPV RNA levels in transfected cells by performing RT-qPCR of CrPV RNA at 72 h.p.t. (Figure 1D). To control for transfection efficiency, the RNA levels were normalized to that within 30 min after transfection. In general, the CrPV RNA levels showed a similar trend as the viral titre measurements. Transfection with ΔVPg1 resulted in similar CrPV RNA accumulation as compared to S2 cells transfected with wild-type CrPV, whereas transfection with ΔVPg1-2 showed significantly reduced viral RNA levels. CrPV RNA levels were at background levels in cells transfected with ΔVPg1-3. In summary, these results strongly suggest that there is a minimum threshold of two VPg copies for productive CrPV infection.

**Figure 1.**
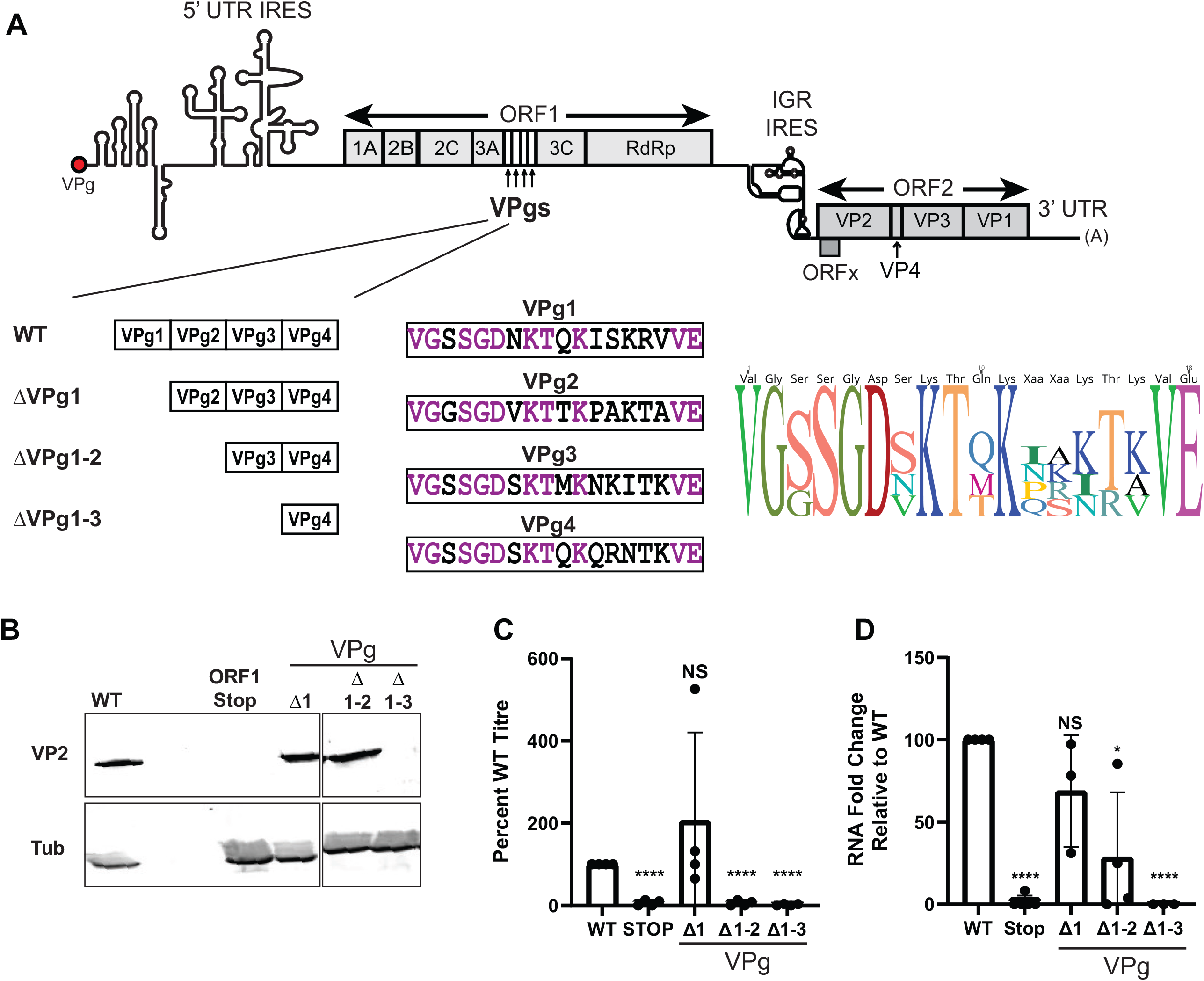
Minimum threshold of VPgs for productive CrPV infection. **(A)** *(Top)* Schematic of the CrPV RNA genome highlighting the arrangement of the four Vpgs *(below, left)* and their amino acid sequences *(below, middle)*. Identical amino acids in all CrPV VPgs are denoted in purple. The VPg deletion mutants within the CrPV-3 infectious clone used in this study are shown (*below, left*). ICE Logo of the 4 CrPV VPgs, created using Geneious (*below, right)*. **(B)** Immunoblots lysates collected from *Drosophila* S2 cells 72 hours post transfection with either WT or mutant CrPV RNA **(C)** Intracellular titres of lysates collected from S2 cells transfected with either WT or mutant CrPV infectious RNA 72 h.p.t. Cells were harvested and titres were measured using VP2 immunofluorescence. (**D)** CrPV RNA levels measured by RT-qPCR of positive-sense RNA compared to RPS6 mRNA at 72 h.p.t. Relative RNA levels (CrPV RNA:RPS6 ratio) were normalized to that at 0.5 h.p.t. for each transfected RNA to control for transfection efficiency and each ratio was normalized to that of the wild-type RNA (100%). Shown are averages from at least three independent experiments with at least three independent technical replicates per experiment +/-SD. *p-value ≤0.05, ****p-value ≤0.0001

### VPg Deletions Attenuate Virus Infection

To investigate infection in a more controlled infectious model system, we propagated wild-type and deletion mutant CrPV from transfected cells. Propagation of virus was possible for all deletion mutant CrPV except ΔVPg1-3, thus reinforcing the importance of the minimal number of VPgs for CrPV infection. After propagating the mutant viruses for three passages, we isolated total RNA, performed RT-PCR, sequenced the VPg region, and confirmed that the deletions were present and stable in the viral genome. We infected naive S2 cells at a MOI of 0.1 and monitored viral yield using an endpoint dilution assay to measure the Median Tissue Culture Infectious Dose (TCID50). Wild-type CrPV infection resulted in peak viral yields by 6 h.p.i. (Figure 2A). By contrast, ΔVPg1 or ΔVPg1-2 CrPV resulted in reduced intracellular viral titres at 6 h.p.i and peaked at 12 h.p.i. (Figure 2A,2C,2D), suggesting a delay in viral infection. For extracellular viral titres, ΔVPg1-2 showed decreased viral titres at both 24 and 48 h.p.i. (Figure 2B,2E,2F), further indicating the importance of VPg copies for optimal CrPV infection.

**Figure 2.**
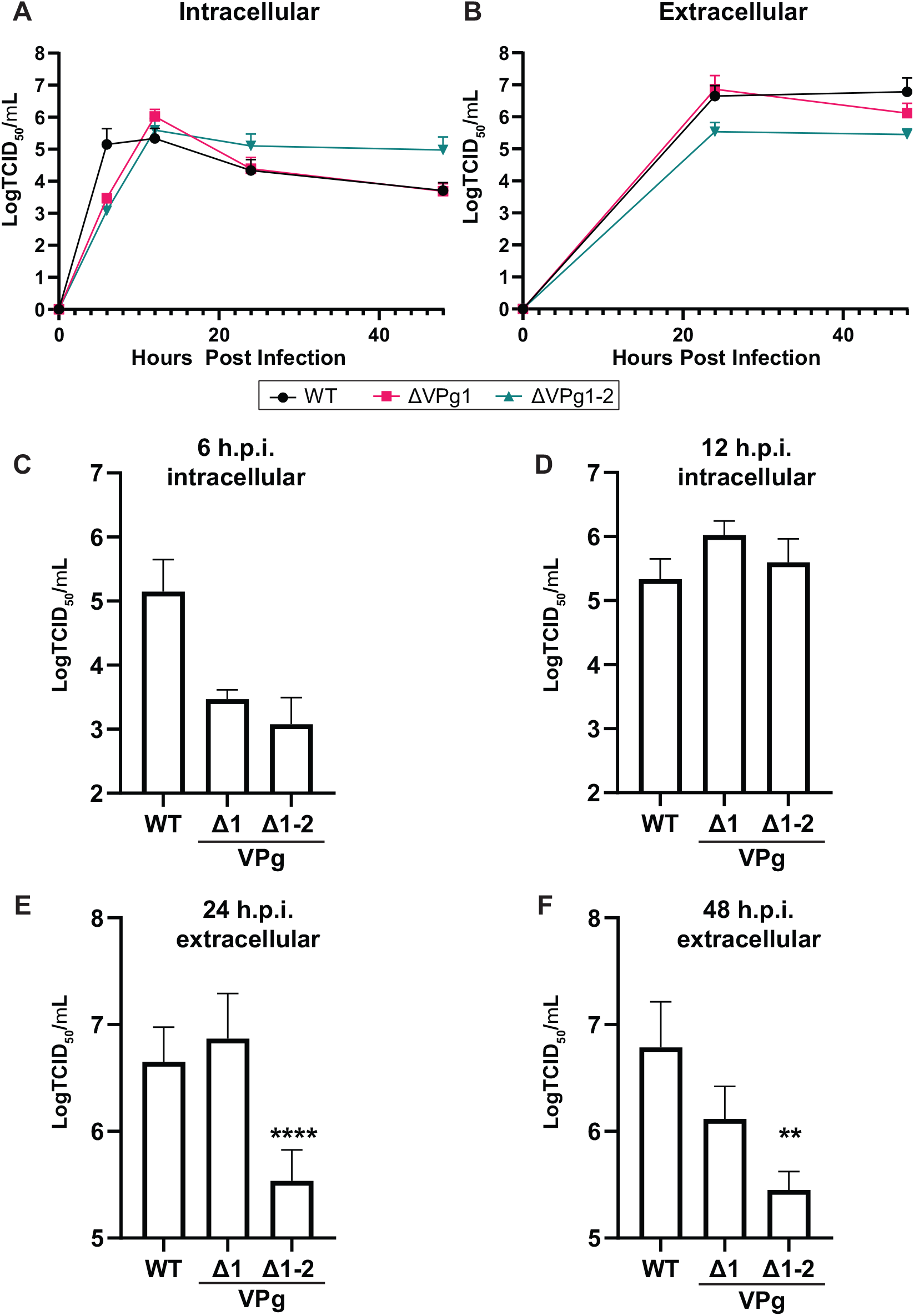
Deletion of VPgs affects CrPV infection. **(A)** Intracellular and **(B)** extracellular titers of WT and mutant CrPV (MOI 0.1) in S2 cells at the indicated times (x-axis) measured by end point dilution (y-axis, log scale). Bar graphs shows intracellular viral titres at **(C)** 6 and **(D)** 12 hours post infection (h.p.i.) and extracellular viral titers at **(E)** 24 and **(F)** 48 h.p.i.. Shown is the TCID50 from at least three independent experiments +/-SD normalized to that of wild-type virus at the corresponding time point for each experiment. **p-value ≤0.01,****p-value ≤0.0001

### CrPV VPg is Linked at the Serine 4 or the Threonine 9 Residue

VPg linkage to the viral RNA is typically through phosphodiester bond acceptor residues, tyrosine, serine, or threonine (50, 51). Picornavirus, calicivirus, and potyvirus RNA genomes are linked to a tyrosine whereas the sobemovirus, Ryegrass mottle virus, utilizes a serine linkage (5). To our knowledge, the only viral genome reported to be linked to a threonine is the sobemovirus, southern bean mosaic virus (5). From the consensus VPg sequence (Figure 1A), several potential phosphodiester bond acceptor residues in CrPV VPgs are possible. In order to determine critical amino acids in CrPV VPgs, we created an alignment of CrPV-like VPgs by aligning previously known (48) and newly discovered (see Materials and Methods) dicistrovirus VPg sequences. Using the consensus CrPV-like VPg sequences, we performed a hidden Markov model search against orthornaviruses to identify RNA viral genomes that contain similar VPgs (52).

We identified 28 unique dicistrovirus genomes including isolates that contain a VPg sequence that passed the search threshold including 11 isolates of drosophila C virus (DCV) and 5 isolates of CrPV (Figure S1A). Alignment of all of these VPgs revealed specific trends, notably an absolute conserved SGDx(K/R)T sequence that is present in every VPg sequence (Figure S1A). Notably, serine 4 and threonine 9 are phosphodiester bond acceptor residues that can be potentially linked to the viral genome. Moreover, there was a high degree of conservation within the N terminal 10 amino acids compared to the C terminus 8 amino acids (Figure S1B). Uniquely the presumed cleavage site differed between different viruses and isolates; E/V, E/I or E/M (Figure S1A). The VPgs shared a 59% pairwise identity and had a high isoelectric point with an enriched basic C terminal end, properties typical of VPg (Figure S1D) (53, 54). After excluding isolates, we identified fourteen unique dicistrovirus genomes using the hidden Markov model (Figure S1C), each of which encoding consecutive tandem VPgs that ranged between two and four copies (Figure S1C). In summary, these analyses identified potential consensus residues that may be critical for VPg function and tandem CrPV-like VPg copies in a subset of dicistrovirus genomes.

To biochemically elucidate the residue linked to the CrPV RNA, we performed mass-spectrometry (MS) analysis on the CrPV VPg using an adapted approach as previously reported (5). Briefly, viral RNA extracted from pelleted CrPV virions was subject to acid-catalyzed hydrolysis using trifluoroacetic acid, which would leave a pUp, pAp, pCp, or pGp on the acceptor residue of VPg. Our proteomic analysis identified VPg2, VPg3, and VPg4 peptides, suggesting that these VPgs are linked to the CrPV RNA genome. We did not identify peptides that overlapped VPg boundaries, suggesting that VPgs are fully processed in CrPV virions. Interestingly, we identified spectra of distinct peptides that contains a pUp modification (+386) on the 9th position threonine of VPg2 (Figure 3A) and on the 4th position serine of VPg4 (Figure 3B) with high confidence (score >100). We also identified a pUp on the serine of VPg2, albeit with a lower score. These results suggested that both the serine 4 and the threonine 9 of CrPV VPg are capable of being uridylated. We did not identify VPg peptides with two pUp modifications, suggesting that uridylation occurs only once per VPg. To our knowledge, this is the first report of a viral RNA linked to multiple residues on a VPg.

**Figure 3.**
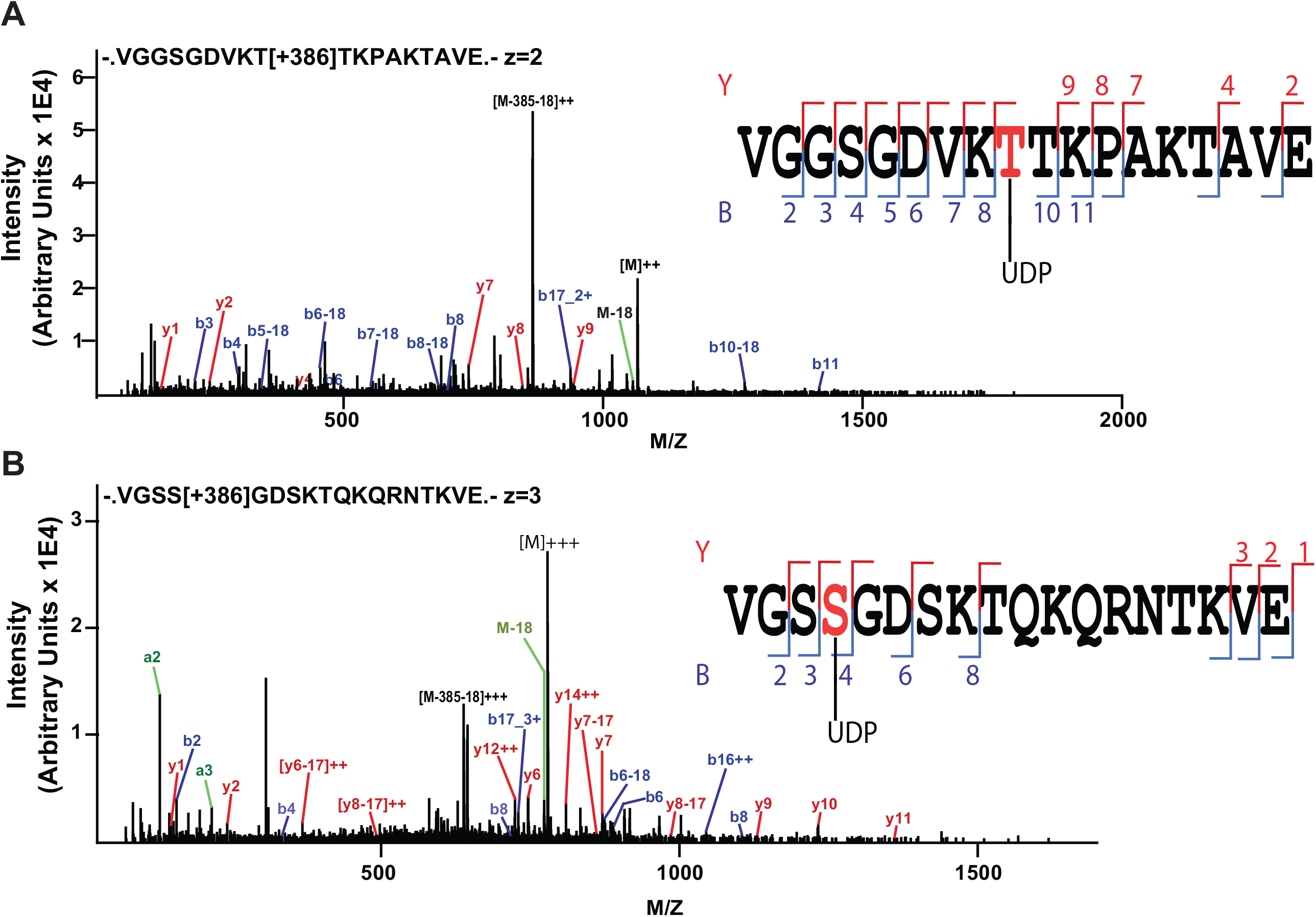
Identification of the VPg linkages by mass spectrometry. **(A)** Fragment spectra from charged VPg2 linked to a uridine via threonine 9. **(B)** Fragment spectra from charged VPg4 linked to a uridine via serine 4. Individual fragment ions are annotated in the spectrum and in the sequence representation. Spectra were viewed and annotated using PMI-Byonic-Viewer (Protein Metrics)

### VPg Serine 4 and Threonine 9 are Essential for CrPV Infection

To determine whether the S4 or T9 of VPgs are important for CrPV infection, we generated mutant CrPV clones containing either S4 to alanine or T9 to alanine in all four VPgs (All S4A or All T9A, respectively) as well as clones containing a single mutation at either S4 or T9 within an individual VPg (Figure 4A). We transfected S2 cells with *in vitro* transcribed wild-type and mutant CrPV RNAs and monitored CrPV infection by immunoblotting for VP2 after transfection (Figure 4B). No VP2 was detected in cells transfected with All S4A or All T9A CrPV RNAs, thus indicating that mutating all S4 or T9 are not viable for CrPV infection (Figure 4B). Furthermore, cells transfected with All S4A or All T9A CrPV RNAs did not show an increase in viral RNAs, thus pointing to a defect in replication (Figure 4C).

**Figure 4.**
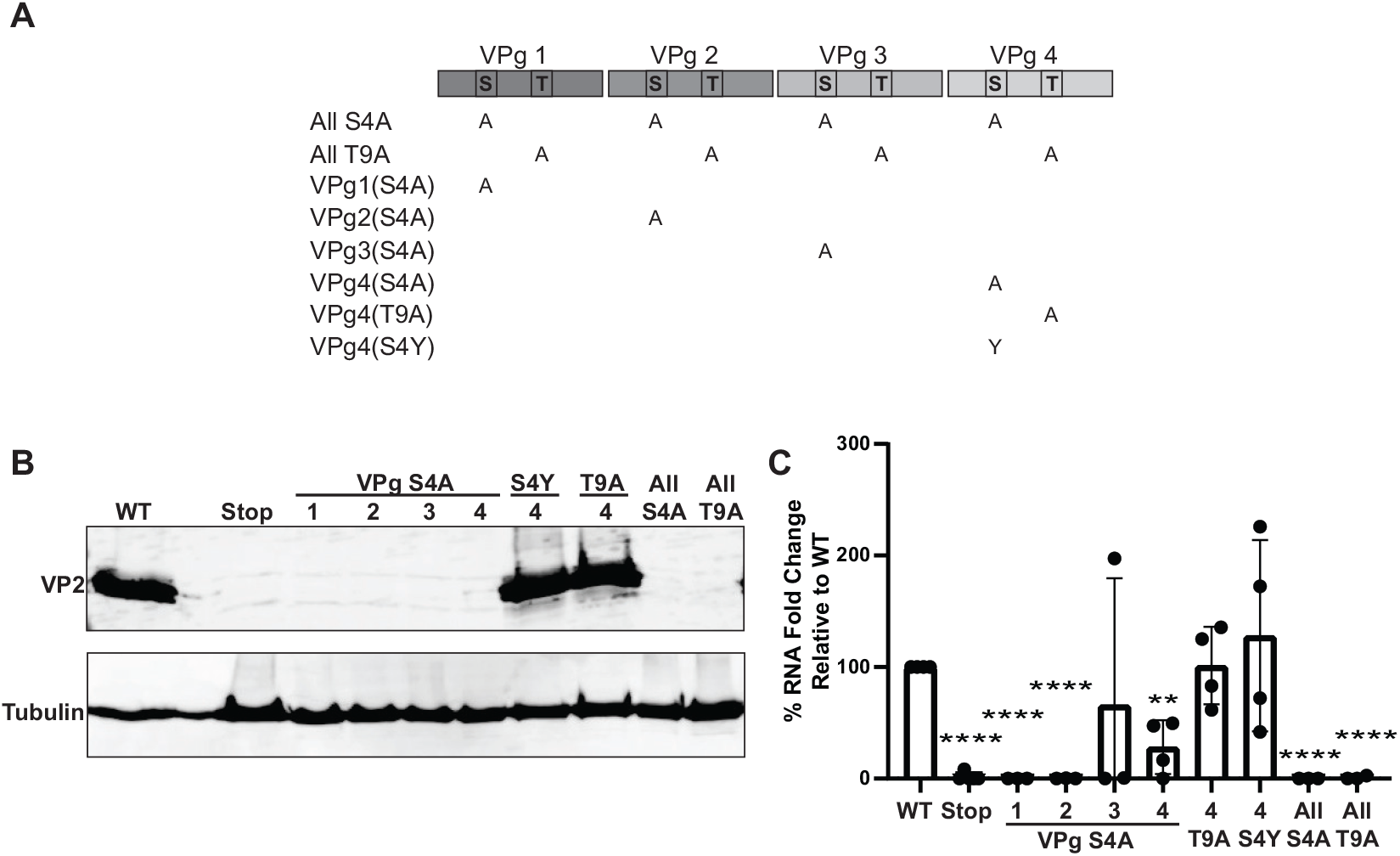
Mutational analysis of VPg on CrPV infection. **(A)** Schematic of S4 or T9 mutations within the VPgs of the CrPV infectious clone. **(B)** VP2 immunoblots of lysates or (**C)** CrPV RNA levels measured by RT-qPCR of positive-sense RNA compared to RPS6 mRNA at 72 h.p.t. Relative RNA levels (CrPV RNA:RPS6 ratio) were normalized to that at 0.5 h.p.t. for each transfected RNA and each ratio was normalized to that of the wild-type RNA (100%). Shown are averages from at least three independent experiments with at least three independent technical replicates per experiment. **p-value ≤0.01, ****p-value ≤0.0001

We next addressed whether specific S4 or T9 within each VPg is important for CrPV infection. Transfection of S2 cells with CrPV RNA containing a mutant VPg4 with a T9A mutation (VPg4(T9A)) resulted in VP2 expression (Figure 4B) and similar levels of RNA accumulation as compared to wild-type CrPV, thus indicating that a single T9A mutation in a VPg does not affect replication. By contrast, transfection of S2 cells with CrPV RNA containing an S4A in any of the VPgs resulted in reduced viral RNA levels and the lack of detctable VP2 expression (Figure 4B,4C), indicating a dominant effect of a single VPg S4A mutation and that the remaining wild-type VPgs cannot compensate.

Through multiple transfection experiments, a fraction of the transfected cells with the single VPg S4A mutant CrPV RNAs showed VP2 expression, suggesting a revertant event. To address this, we extracted total RNA from a subset of transfected cells that showed VP2 expression, performed RT-PCR, and sequenced the VPg region. The results showed several sequences that deviated from the S4A mutation including reversion of the mutant alanine 4 back to serine or deletion of multiple VPgs. These results indicated that the S4A mutation in a single VPg is relatively unstable and highlights the importance of the S4 residue of VPg for replication.

To determine the specificity of the VPg S4 for viral replication, we generated a CrPV clone containing mutation of S4 to tyrosine of VPg4, which has the potential to be a phosphodiester bond linking acceptor. Transfection of the CrPV VPg4(S4Y) RNA resulted in VP2 expression and in detectable replicative RNA levels comparable to wild-type CrPV (Figure 4B, 4C). These results suggested that mutating S4 to tyrosine of a VPg does not affect virus infection. In summary, these results all point to S4 as the primary linking amino acid to the CrPV RNA that is essential for CrPV replication.

### Polyprotein processing of mutant CrPV VPg in vitro

Given that VPg3 mutations disrupt polyprotein processing in FMDV (29), we examined whether deletion of VPgs or the S4A VPg mutants results in a defect in polyprotein processing. To address this, we monitored translation and polyprotein processing *in vitro* by incubating wild-type and mutant CrPV VPg RNA in Sf-21 translation extracts. For all experiments, equal amounts of *in vitro* transcribed RNA were verified (Figure 5B). Incubation of wild-type CrPV RNA in Sf-21 extracts resulted in translation of non-structural and structural proteins and processing of the polyproteins as shown previously (Figure 5) (49). Notably, several precursor proteins such as the full-length ORF2 and mature proteins, VP2/VP3 and 1A, were detected. A mutant CrPV RNA containing a stop codon within ORF1, which inhibits 3C protease expression, led to only unprocessed ORF2 expression (49). For the single VPg deletion mutants (ΔVPg1), a ∼63 KDa protein is detected which is ∼2 KDa smaller than the full-length precursor 3A-VPg-3C. Progressive deletion of VPgs (ΔVPg1-2 and ΔVPg1-3) resulted in a progressive smaller precursor 3A-VPg-3C protein (Figure 5B). Besides these differences, the pattern of detectable proteins suggested that there are no major defects in polyprotein processing when VPgs are deleted as compared to that with the CrPV RNA in this *in vitro* system. However, although polyprotein processing was in general similar to that of the wild-type CrPV, subtle differences were observed with the S4A mutation in a single VPg. For example, a ∼60-65 kDa band is reproducibly detected in the reactions with S4A VPg3 but not with the other S4A VPg mutants (asterisk, Figure 5B). Furthermore, detection of a ∼25 kDa protein, presumably the 3AB precursor protein, is not detected in a fraction of experiments with S4A VPg3 and VPg4 but present in S4A VPg1 and VPg2 reactions. Detection of the 25 kDa protein was sporadic between experiments suggesting a minor processing defect by the mutant VPg. In summary, these results indicated that although there were no other major defects in processing, there may be specific albeit subtle alternative processing events attributed to a subset of single S4A VPg RNAs *in vitro*.

**Figure 5.**
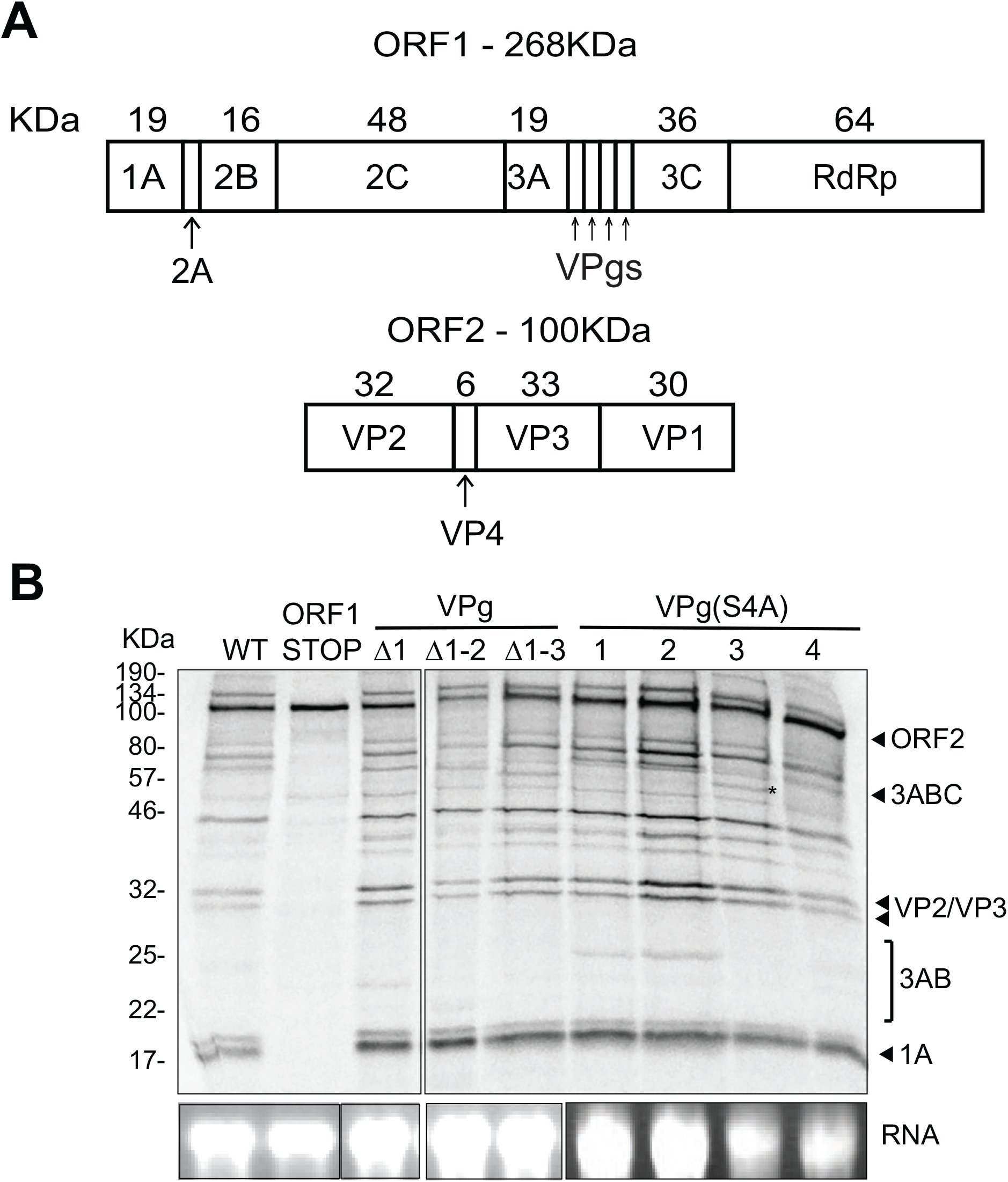
Translation and polyprotein processing of CrPV VPg mutant RNAs *in vitro*. **(A)** Schematic of expected protein sizes of proteins synthesized from ORF1 (top) or ORF2 (bottom) of CrPV **(B)** Audoradiograph of *in vitro* translated CrPV WT or mutant Shown is a representative gel from at least three independent experiments. (*) asterisk denotes a specific altered processed viral protein. (*below*) Gel analysis of *in vitro* transcribed RNA (0.5 µg) detected by ethidium bromide.

### Multiple VPgs Compensate for IRES Translation Mechanisms

Dicistroviruses utilize a viral translation strategy whereby the distinct IRES translation mechanisms result in molar excess expression of ORF2 over that of ORF1 (46, 55). The 5’ UTR IRES that drives ORF1 expression is relatively weaker compared to that of the IGR IRES that directs ORF2 expression. Nakashima *et al*. proposed that the multiple VPgs in dicistrovirus genomes compensate for the differences in translation of ORF1 and ORF2 (48). To address this hypothesis, we monitored the translation of the 5’ UTR and IGR IRESs from dicistrovirus genomes that differed in the number of VPgs. We identified the presence of multiple VPgs using a RADAR search (56). Specifically, we chose the following dicistrovirus genomes with varying numbers of VPgs: Triatoma virus (TrV; NC_003783) with 2 VPgs, Cricket paralysis virus (CrPV; NC_003924) with 4 VPgs, Solenopsis invicta virus 1 (SINV-1; NC_006559) with 6 VPgs, and Macrobachium rosenbergii taihu virus (MrTV; NC_018570) with 7 VPgs. Towards this, we generated “minigenomes” of dicistroviruses whereby we replaced the ORF1 and ORF2 with renilla and firefly luciferase, respectively, to produce a reporter system to monitor translation (Figure 6A).

**Figure 6.**
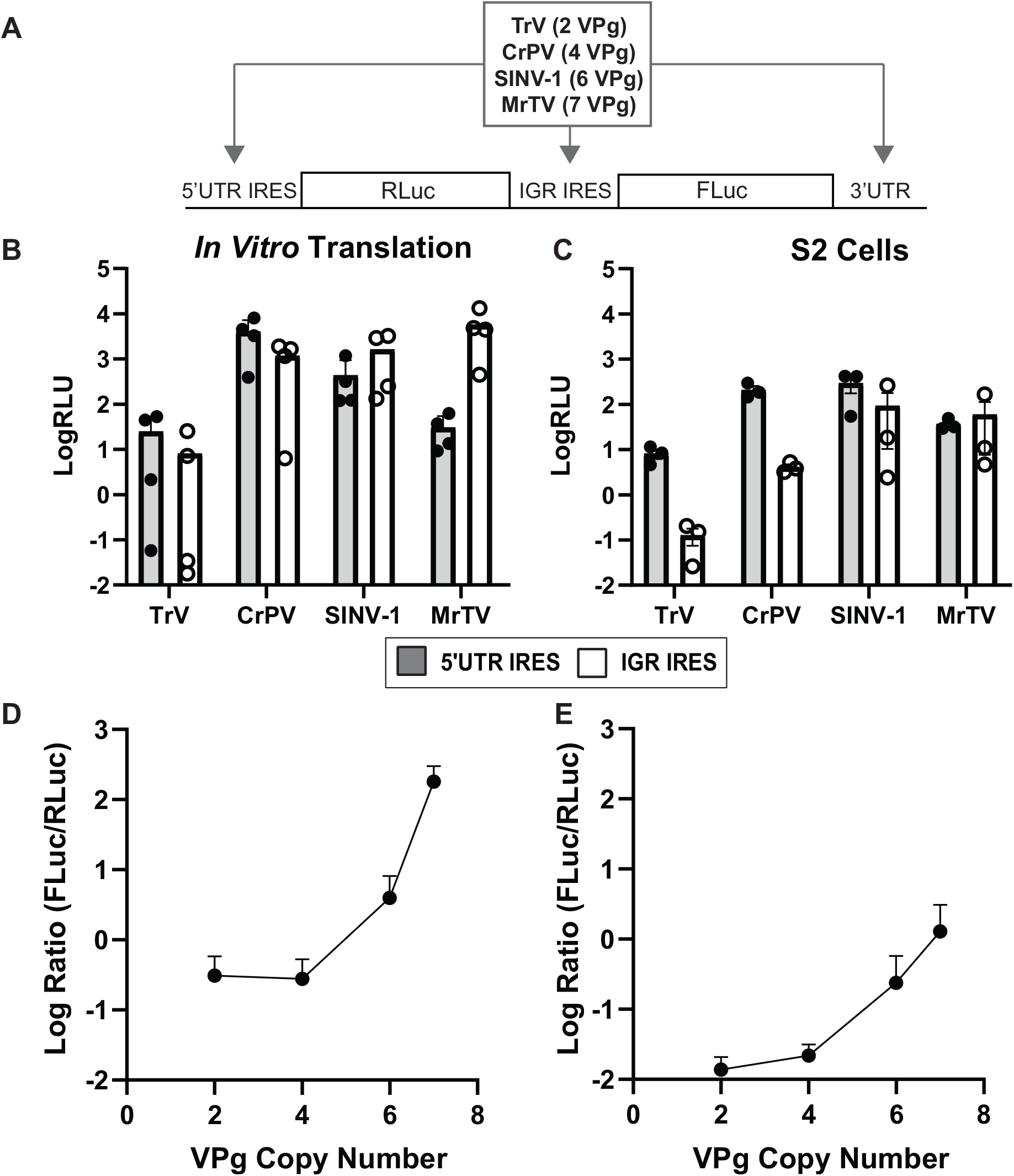
Dicistrovirus IRES translation correlates with VPg copy number. **(A)** Schematic of minigenome reporters that contain 5’ UTR, IGR and 3’UTR sequences from four dicistrovirus genomes. Dicistrovirus genomes were selected based on the number of VPgs. The 5’ UTR and IGR IRESs direct renilla (RLuc) and firefly (FLuc) luciferase translation, respectively. In vitro transcribed minigenome RNAs were either **(B)** incubated in Sf21 translation extracts or **(D)** transfected into CrPV-infected cells. Renilla and firefly luciferase were monitored after 1.5 hours in translation extracts or at 6 hours post infection. Shown in **(C)** and **(E)** are the ratio of FLuc:RLuc luciferase activities (y-axis) versus the number of VPgs in the dicistrovirus genomes (x-axis). Shown are averages +/-s.d. from at least three independent experiments.

We incubated *in vitro* transcribed “minigenome” RNAs in Sf-21 insect cell extracts and monitored luciferase activities (Figure 6B, 6C). All minigenome RNA led to luciferase activities to varying degrees, which is indicative of active 5’ UTR and IGR IRES translation (Figure 6B, 6C). We calculated the relative ratio of Fluc:RLuc, with the prediction that there would be a correlation with VPg copy numbers for each genome. This analysis, in general, revealed that the FLuc:RLuc ratios increased exponentially with the number of VPg copies, with the highest ratio in the MrTV minigenome.

However, the trend was not clear for the viral genomes with fewer VPgs. For instance, the TrV and CrPV minigenomes showed similar FLuc:RLuc ratios although the ratios for these were much lower compared to that of the minigenomes under MrTV.

We next examined the translation of these minigenomes in a more physiological condition under CrPV infection in S2 cells. Specifically, S2 cells were transfected with the minigenome RNAs (one hour prior) and then infecting cells with CrPV and luciferase activities monitored at six hours post-infection. For all minigenomes, 5’ UTR and IGR IRES-mediated luciferase activities increased in infected cells compared to uninfected cells (46) (Figure 6C, 6E). Similar to the *in vitro* translation results, the MrTV minigenome showed the highest FLuc:RLuc ratio in CrPV infected cells followed by SINV-1, CrPV, and TrV. Plotting the FLuc:RLuc ratio against the number of VPgs revealed a strong logarithmic correlation (R>0.9) (Figure 6E), thus supporting the idea that VPg copy numbers within the dicistrovirus genome compensate for the lower 5’ UTR IRES translational activity compared to that of the IGR IRES.

## DISCUSSION

Due to their size and error prone RdRP, genetic duplications are not maintained and thus are rare events in RNA viruses (57). There must be strong selection pressure for a gene duplication in a viral genome and therefore must be essential for virus infection. However, the mechanisms underlying this are not fully understood. In this study, we provide insights into the function of duplicated VPgs in a viral genome. Using the dicistrovirus CrPV as a model, we determined that there is a minimal number of VPgs for productive infection. We identified two residues, S4 and T9, that are linked to the CrPV RNA genome and through mutational analysis, we showed that S4 is a critical residue for viral infection. To our knowledge, this is the first report of a viral RNA that is capable of linking to two residues on VPg. Finally, we demonstrated that VPg copies compensate for the distinct IRES translational activities.

In FMDV infections, multiple VPgs are required for optimal replication, however, deletion of two of the three VPgs was still viable (58). In another study, deletion of the first two VPgs did not support FMDV infection in porcine cells but was infectious in hamster and bovine cells, thus pointing to a role of the VPg proteins in specifying virus host range (59). In the case of CrPV, a minimal number of VPgs (two) are required for productive replication and infection. Although there were no major differences in CrPV polyprotein processing *in vitro* (Figure 5), studies on FMDV showed that deletion of VPg3 had an effect on polyprotein processing (25, 29). It remains to be investigated whether depletion of other VPgs in CrPV affects a distinct step in the viral lifecycle.

Unlike the single large ORF in FMDV, the dicistroviruses contain two main ORFs that are controlled by distinct IRES mechanisms. Nakashima *et al*. proposed that multiple VPgs in dicistroviruses compensate for the relative translational activity of ORF1 compared to that of ORF2 (48). In support of this, we demonstrated that the number of VPgs within dicistrovirus genomes correlates with the ratio of IGR IRES:5’ UTR IRES translational activities. As VPg is required for replication for each viral RNA, it is imperative that there is sufficient VPg levels for optimal viral replication. Thus, multiple copies of VPg within ORF1, which is expressed much lower relative to that of ORF2, are needed to compensate for the number of VPgs expressed for optimal replication. This finding is further supported by the lower titre following infection of virus with only two VPgs (Figure 2).

Our mass spectrometry approach identified two residues within VPgs that can be linked to the RNA genome. An outstanding question is how can a CrPV VPg be uridylated at two distinct residues? Picornavirus VPgs typically contain positively charged amino acids that allow binding to the negatively charged binding groove of RdRP (60). Although picornavirus RdRPs and VPgs have similar structures, structural studies of CVB3, FMDV, and EV71 RdRP:VPg complexes revealed distinct binding sites, suggesting that each virus uses different uridylation mechanisms (61). Moreover, it has been proposed that uridylation of EV71 VPg is mediated by a two-step mechanism requiring two EV71 RdRPs binding to VPg (62).

Our mutagenesis studies showed that the S4 and T9 are not functionally redundant for viral replication. Mutating S4 or T9 in all VPgs did not yield productive infection (Figure 4). However, mutating T9 to alanine in two VPgs led to productive virus infection (data not shown). The exact role of the genome linkage to T9 is not clear. It is possible that an alternative uridylation site for VPgs is required at a specific step in infection to ensure promotion of replication. Our results point to S4 as the linking amino acid of CrPV VPg to the viral genome. The recovery of revertants containing alanine to serine residues in VPg supports the importance of the S4 in CrPV replication.

Interestingly, mutating this same residue to a tyrosine instead of an alanine recovered replication similar to that of wild-type infection (Figure 4). Our findings that an S4A mutation in a single VPg abolished infection but not when a VPg is deleted suggests a dominant-negative effect. It is probable that the assembly of the replication complex is rate-limiting and that a single VPg containing the S4A blocks an assembly step or replicase activity, possibly at the CRE element and/or the priming step. This finding is similar to that observed in FMDV whereby mutating the linking tyrosine of VPg1 to a phenylalanine resulted in a larger reduction in titre than deleting that same VPg (25). Trans-acting dominant effects of VPg mutants have also been observed with poliovirus infection (63), thus suggesting a rate-limiting step in replication. Alternatively, the dominant effects by a single S4A VPg mutant may affect other steps in the viral lifecycle such as polyprotein processing. Our results showed that a subset of an S4A mutant in a single VPg led to subtle polyprotein processing defects *in vitro* (Figure 5). The exact nature of these unprocessed or defective processing events is not clear but does hint of multifunctional roles of the CrPV VPgs in replication and polyprotein processing. Indeed, studies on FMDV have shown that the C-terminal amino acids of VPg3 specifically at the VP3-3C boundary are important determinants in the rate of processing (29). It will be of interest to examine how N-terminal mutations within each CrPV VPg (i.e. S4A) affects processing rates at the VPg-VPg and VpG-3C boundaries. Further investigation using *trans*-complementation experiments, reconstituted replication systems and monitoring polyprotein processing in infected cells should shed light into the roles of the CrPV VPg mutants.

Positive strand RNA viruses have limited coding capacity, thus the presence and the strong maintenance multiple copies of VPgs in a viral family indicates a functional importance of VPgs in viral replication. Tandem VPgs have roles in compensation effects on viral translation (this study) and polyprotein processing (29). A number of Dicistrovirus genomes appear to have multiple VPgs that differ in sequence and potentially diversity in residues that can be linked to viral RNA (48). Thus dicistro-like genomes are a rich source of viral models to examine the mechanistic role(s) of tandem VPg-like sequences during viral infection.

## Acknowledgements

We thank the Jan lab for critical analysis of the study and editing of the manuscript. We would also like to thank Morris Young for assistance in cloning a subset of constructs. This study was supported by a CIHR operating grant (PJT-178342) and an NSERC Discovery grant (RGPIN-2017-04515) to EJ, an NSERC DG and Genome Canada (Project 264PRO) grant to LF, and a NSERC PGS D scholarship to RW.

## Materials and Methods

### Cell Culture and Virion Isolation

Drosophila Schneider line 2 (S2) cells were maintained and passaged in schneider’s insect medium supplemented with 10% fetal bovine serum. Propagation of CrPV in Drosophila S2 cells was performed as previously described (49). Briefly, S2 cells were infected with CrPV for a minimum of 2 days and then lysed in 0.5% IGEPAL CA-630 (NP-40) and 0.1% 2-mercaptoethanol on ice for 10 minutes. Cell debris was cleared at 13 800 RCF for 15 minutes at 4°C. The supernatant was then centrifuged at 141 000 RCF for 2 hours. The viral pellet was resuspended in PBS and the viral stock aliquoted and stored at -80C. Virus was then tittered by end-point dilution.

### Plasmids and Transfections

Deletion mutations were introduced into the CrPV infectious clone (CrPV-3) by Gibson assembly (Gibson et al., 2009). Mutations were generated using quick-change PCR as previously described (Bachman, 2013). For the minigenome constructs, the SINV-1 minigenome was generated (NC_006559) (Twist Bioscience (San Francisco)). The T7 promoter-containing clone (backbone pTwist Amp High copy) encodes the renilla and firefly luciferase open reading frames and the following: SINV-1 5’ UTR is flanked between HindIII and KpnI site, the SINV-1 IGR IRES is flanked between EcoRI and BamHI, and the the SINV-1 3’UTR is between XbaI and SpeI. The 5’ UTRs, IGR IRESs and 3’UTRs of other dicistroviruses (Twist Biosciences, genewiz for MrTV) were cloned into the corresponding restriction sites to generate minigenomes. All clones were sequence confirmed. Plasmids containing the CrPV or the minigenomes were linearized with Ecl136ii (Thermofisher) or Eco53KI (NEB). The CrPV minigenome was linearized using BamHI. RNA was transcribed using a T7 RNA polymerase reaction and subsequently DNAse-treated for one hour followed by purification using a RNeasy kit (Qiagen) as per manufacturer’s instructions. The integrity and purity of the RNA was confirmed on a 1% denaturing formaldehyde agarose gel.

### *In Vitro* Translation

*In vitro* transcribed CrPV RNA was incubated in Sf-21 cell extract (Promega) in the presence of [^35^S]-methionine-cysteine for 2 hours at 30°C. Equal volumes of reaction mixtures were loaded on SDS-PAGE gels. Gels were dried, and radioactive bands were monitored by phosphoimager analysis (General Electric)

### Viral Titres

Drosophila S2 cells transfected with mutant CrPV RNA or mutant CrPV virus were collected and lysed by a 3X freeze-thaw. Following lysis, debris was spun down at 13200 rpm for 30 minutes at 4°C. TCID_50_: A 96 well plate was seeded with 3E5 S2 cells per well. The lysed materials were then serially diluted, and 8 wells were given each dilution. The cells were then incubated for 5 days at 25°C and the presence of cytopathic effects was determined for each well and the TCID_50_ was calculated. CrPV VP2 Immunofluorescence: immunofluorescence was performed as previously described (49).

### Western Blotting

Equal amounts of S2 protein lysates were resolved on a 15% SDS-page gel then transferred to a polyvinylidene difluoride immobilon-FL membrane. Membranes were blocked for 30 minutes at room temperature with 5% skim milk in tris-buffered saline plus tween-20 (TBST). The blots were incubated for 1 hour at room temperature with CrPV (VP2) ORF2 rabbit polyclonal (1:1000) antibody as described (49) and tubulin antibody (1:1000) (DSHB E7). Membranes were washed three times with TBST and incubated with IRDdye 800CW goat anti-rabbit IgG (1:10000; Li-cor biosciences) and then detected by Odyssey imager.

### Mass-Spectrometry

After cell lysis, CrPV virions were spun down at 141000 RCF. The RNA was extracted from virions using a RNAeasy miniprep kit (Qiagen) and the RNA was hydrolyzed using trifluoroacetic acid to a final concentration of 10%. Analysis was performed using Peptide samples were purified by solid phase extraction on C-18 stage tips (64–66). Purified peptides were analyzed using a quadrupole – time of flight mass spectrometer (Impact II; Bruker Daltonics) on-line coupled to an Easy nano LC 1000 HPLC (ThermoFisher Scientific) using a Captive spray nanospray ionization source (Bruker Daltonics) including a 75-μm-inner diameter, 40 cm long fused silica analytical column with an integrated spray tip (6 – 8 μm-diameter opening, pulled on a P-2000 laser puller from Sutter Instruments). The analytical column was packed with 1.9 μm-diameter Reprosil-Pur C-18-AQ beads (Dr. Maisch) and it was heated to 50°C using tape heater (SRMU020124 and in house build temperature controller). Buffer A consisted of 0.1% aqueous formic acid and 2 % acetonitrile in water, and buffer B consisted of 0.1% formic acid in 90 % acetonitrile. Samples were resuspended in buffer A and loaded with the same buffer. For 60 min run the gradient was from 5% B to 30% B over 60 min, then to 100% B over 2 min, held at 100% B for 13 min. Before each run the analytical column was conditioned with 4 μL of buffer A and the sample loading was set at 10 μL (for samples up to 3 μL volume). The LC thermostat temperature was set at 7°C. The Captive Spray Tip holder was modified similarly to an already described procedure (67) the fused silica spray capillary was removed (together with the tubing which holds it) to reduce the dead volume, and the analytical column tip was fitted in the Bruker spray tip holder using a piece of 1/16” x 0.015 PEEK tubing (IDEX), an 1/16” metal two way connector and a 16-004 Vespel ferrule. The sample was loaded on the trap column at 950 Bar and the analysis was performed at 0.35 μL/min flow rate. Impact II was run with OTOF Control v. 4.1 (Bruker). LC and MS were controlled with HyStar 4.1 (4.1.21.2, Bruker). The Impact II was set to acquire in a data-dependent auto-MS/MS mode with inactive focus fragmenting the 20 most abundant ions (one at the time at 18 Hz rate) after each full-range scan from m/z 200 Th to m/z 2000 Th (at 5 Hz rate). The isolation window for MS/MS was 2 to 3 Th depending on parent ion mass to charge ratio and the collision energy ranged from 23 to 65 eV depending on ion mass and charge (67). Parent ions were then excluded from MS/MS for the next 0.3 min and reconsidered if their intensity increased more than 5 times. Singly charged ions were excluded since in ESI mode peptides usually carry multiple charges. Strict active exclusion was applied. Mass accuracy: error of mass measurement is typically within 5 ppm and is not allowed to exceed 10 ppm. The nano ESI source was operated at 1900 V capillary voltage, 0.25 Bar nanoBuster pressure with methanol in the nanoBooster, 3 L/min drying gas and 150°C drying temperature. Mass spectrometry data were processed using Byonic (Protein Metrics) searching against the expected peptides with variable modifications phosphodiester (pUp, pAp, pCp or pGp) at serine or threonine.

### VPg Sequence Identification

Using the CrPV 2C and 3C protein sequences as queries (Accession: Q9IJX4), the Dicistroviridae protein sequences available at NCBI were searched with blastp (68). The blastp hits were then processed in order to extract the amino acid sequences between 2C and 3C plus an additional 100 amino acids. The resulting sequences were then searched with locally installed versions of HHrepID (69), XSTREAM (70) and TRUST (71) to identify potential tandem repeats. The tandem repeat hits were then manually curated and sorted by length and initial amino acid to obtain high-quality representative VPgs. A multiple alignment of CrPV like VPgs was constructed with Clustal Omega (72) which was then used to build a hidden Markov model with the HHMER toolkit (73). All RefSeq viral sequences downloaded from NCBI were then searched using those hidden Markov models associated with high-quality VPgs.

Rapid automatic detection and alignment of repeats in protein sequences was performed by downloading the “dicistroviridae” sequences from NCBI virus (74) and entering into the RADAR program (56). The VPg like sequences were copied into a FASTA file for use in alignments.

### Luciferase Assay

For *in vitro* translation assays, 1 µg of the RNA was placed in Sf21 cell extract and a reaction was performed for 1.5 hours at 30°C (75). 1 µL of this reaction was diluted in 100uL of passive lysis buffer (Promega). Following this, a 10 µL aliquot was incubated with 50 µL of LARII (Promega) and read using a TD20/20 luminometer (Turner). Then 50 µL of Stop-Glo (Promega) was added to read the renilla luciferase activity.

For translation assays in S2 cells, minigenomes were transfected one hour prior to infection with CrPV. CrPV infection was performed at MOI 20 for 6 hours and then cells were lysed in passive lysis buffer (Promega) and luciferase activities measured by a dual-luciferase assay (Promega).

### RT-qPCR

3 µg of RNA was diluted in Opti-MEM with 2.5uL of lipofectamine 2000 (Life Technologies). Half of the RNA-lipofectamine mixture was added to 1.5X10^6^ S2 cells on a 12 well plate. Transfection mixture was allowed to absorb for 15 minutes before being washed off and 1 mL of schneider’s insect medium supplemented with 10% fetal bovine serum was added. At corresponding timepoints (30 minutes and 72 hours), cells were pipetted off the plate, spun down and washed once with 1X PBS. The RNA was then extracted using a Monarch total RNA miniprep kit (NEB) as per manufacturer’s instructions. RNA was then diluted to 20 ng/uL and 20-60 ng of RNA was used for qPCR using Luna universal one-step RT-qPCR kit (NEB) as per manufacture’s instructions. Primers used for detection were CrPV: 5’-CAGTGCCTTACATTGCCA-3’ and 5’AACTTCTACTCGCACTATTC-3’ Rps6: 5’-CGATATCCTCGGTGACGAGT-3’ and 5’-CCCTTCTTCAAGACGACCAG-3’ All qPCR data was first normalized to RPS6 for the specific sample at the specific time point. CrPV:RPS6 RNA ratio was normalized to the ΔCt 0.5 hours post transfection for each RNA where ΔΔCt is 2^-(ΔCt_72hours_-ΔCt_0.5hours_). The ΔΔCt was normalized to the wild-type CrPV:RPS6 radio (presented as 100%).

## FIGURE LEGENDS

**Figure S1. Dicistroviruses Contain Multiple VPgs. (A)** Alignment of VPg sequences found from hidden Markov search of CrPV like dicistroviruses. Accession number is labelled for each VPg and amino acid numbers. Black indicates full conservation; grey indicates partial conservation. **(B)** ICE logo of alignment provided on the right. Full conservation of SGDx(K/R)T can be seen at 4^th^ position **(C)** Pie chart showing individual viruses that were found and the amount of VPgs found. 3 unique viruses had 2 VPgs, 8 had 3VPgs and 3 unique viruses had 4VPgs **(D)** Average values for all the VPgs found using the HMM search against the CrPV like VPgs. The VPgs are 59% conserved or 33% identical with a molecular weight of 1.869 and an isoelectric point of 10.054.

